## Supplemental Figures for "Energy Coupling and Stoichiometry of Zn^2+^/H^+^ Antiport by the Cation Diffusion Facilitator YiiP"

### Supplemental Figure Legends

#### Suppl. Figure 1. Determination of WT and D51A structures by cryo-EM.

(A,B) SDS-PAGE, left, and elution profile from SEC purification, right. Molecular weight standards are in the first lane at 116, 66, 45, 35, 25, 18.4, 14.4 kDa, with the YiiP monomer running as a single band at 32.5 kDa. (C,D) The cryo-EM workflow starts with correction of beam-induced motion and contrast transfer function (CTF) of movies, followed by template-based particle picking. The particle set was subjected to two rounds of 2D classification and ab initio structure determinations to remove false positives. (E,F) Hetero-refinement jobs were used to look for multiple conformations and to maximize the homogeneity of particles for a final non-uniform refinement job, in this case with C2 symmetry. (G,H) The final structure was characterized by local resolution, represented by the colored surface shown from two orthogonal directions, and the Fourier Shell Correlation.

#### Suppl. Figure 2. Density at $Zn^{2+}$ sites in the cryo-EM maps.

An overview of each cryo-EM structure is shown on the left, followed by densities at sites A, B and C, respectively. Contour levels used to display the various maps were as follows. WT: 6 $\sigma$ for site A, 7 $\sigma$ for sites B and C, D51A: 5.1 $\sigma$ for site A, 7 $\sigma$ for sites B and C, D287A, 6.5 $\sigma$ for sites A and C, 5.5 $\sigma$ for site B, D70A_asym: 9 $\sigma$, D70A_sym: 9 $\sigma$, D287A/H263A: 10 $\sigma$. For clarity, side-chains are shown only for selected residues. In D70A_asym, the TM2/TM3 loop forms a $Zn^{2+}$-free association with the CTD (H1 and C-terminus of M6) in one protomer (panel H) and is disordered in the other protomer (not shown). For D70A_sym, site B is disordered in both protomers and therefore not shown (panel K). In D287A/H263A, one of the TM2/TM3 loops associates with CTD's from opposing protomers (CTDb, CTDc and TMDd refer to chains B, C and D in panel Q, respectively). The TM2/TM3 loops are also shown in Suppl. Fig. 5.

#### Suppl. Figure 3. Determination of D70A structures by cryo-EM

(A) SDS-PAGE and elution profile for SEC purification; molecular weight markers are at 116, 66, 45, 35, 25, 18.4, 14.4 kDa. The workflow for image processing was complex, involving multiple attempts to segregate distinct classes via ab initio and hetero-refinement jobs. This chart is a summary of the final, productive steps leading to structures presented. (B) Initial steps included correction of beam-induced motion and CTF followed by template-based particle picking, rounds of 2D classification and ab initio structure determination. (C) Hetero-refinement was used to segregate two major classes: class 1 comprising the expected dimer and class 2 comprising a dimer of dimers. The dimer of dimers involved interactions between Fab domains and thus did not appear to affect the conformations adopted by YiiP. (D) Particles from class 1 were subjected to further hetero-refinement to segregate into an additional two classes: one with symmetric TMD's (purple) and the other with asymmetric TMD's (blue) as described in the text. (E) Particles from class 2 (dimer of dimers) were exported to RELION where symmetry expansion and signal subtraction was used to generate a larger set comprising isolated dimers. (F) These symmetry-expanded particles were grouped together with class 1 particles for hetero-refinement using reference volumes with symmetric and asymmetric TMD's as well as a junk collector (not shown). (G, H) The resulting final classes were then used for non-uniform refinement to produce D70A_sym and D70A_asym structures, each colored according to local resolution and shown from two orthogonal directions.

##### Suppl. Figure 4. Determination of D287A and D287A/H263A structures by cryo-EM

(A,B) SDS-PAGE and elution profiles from SEC purifications. Images come from a single gel with the molecular weight markers (116, 66, 45, 35, 25, 18.4, 14.4 kDa). Position and presence of the double peak is consistent with the higher order oligomerization seen during image analysis. (C,D) Corrections for motion and CTF were followed by two rounds of 2D classification and ab initio structure determination. (E,F) Hetero-refinement was used to select a homogeneous set of particles for non-uniform refinement. (G,H) Final structures are characterized by FSC and by local resolution, illustrated by the plots and by surface coloring, respectively.

##### Suppl. Fig. 5. Interactions between the TM2/TM3 loop and the CTD.

(A) In the WT structure, the TM2/TM3 loop (blue) contacts the TM6/CTD linker from the opposing protomer with proximity of D72 and R210. (B) In the D70A_asym structure, the $Zn^{2+}$-free TM2/TM3 loop extends to interact with the H1 helix in the CTD of the opposing protomer. (C) In the D287A/H263A structure, the TM2/TM3 loop extends towards CTD's of two different protomers (e.g., TM2/TM3 loop from chain B inserts between CTD's of chains A and D). (D) In the other monomer of D287A/H263A, the TM2/TM3 loop interacts with the TM6/CTD linker of an opposing chain (e.g., the loop from chain A interacts with the linker from chain B). In these chains, the TM6/CTD linker has refolded into one, long continuous helix.

##### Suppl. Fig. 6. Measuring $Zn^{2+}$ affinity by MST

(A-D) SEC elution profiles for YiiP constructs. WT protein is included for comparison but was not analyzed by MST. Elution volumes of the peak are indicated on each plot, which are almost identical for all constructs. (E-G) Raw data from MST for the titrations at pH 7. The blue rectangles correspond to the time window used for initial fluorescence levels and the red rectangles correspond to the time window used for the thermophoresis rates. Each plot includes 48 traces that represent data from 16 $Zn^{2+}$ concentrations used for the titration, each of which were measured in triplicate. (H) SEC profiles of the D287A mutant in the presence and absence of Fab. The shift in position of the main peak in the presence of Fab is consistent with a complex between the homodimer and two Fab's and the secondary peak at 9.9 ml is consistent with the dimer of dimers (4 YiiP + 4 Fab) seen during cryo-EM process (Suppl. Fig. 4). (I) SEC profiles of the D287A/H263A mutant in the presence and absence of Fab. Although the position of the peak in the absence of Fab is consistent with the YiiP homodimer seen with other mutants, the position of the main peak in the presence of Fab is consistent with a dimer of dimers and the secondary peak with even higher order oligomers.

##### Suppl. Fig. 7. Walk of replica simulations in CpHMD pH-replica exchange (REX) through pH space.

Each panel shows how one replica simulation (Rep:0 to Rep:29) changes over the course of the simulation (Frame number) its current pH state, as indicated by the "pH replica" on the ordinate. The pH replicas range from pH 1.5 (pH replica 0) to 11.5 (pH replica 29). Multiple replicas sample most of the pH range and many move across the especially relevant range between pH replica 9 (pH 4) and pH replica 22 (pH 9), indicating that the REX procedure samples pH space well.

#### Suppl. Fig. 8. Convergence of the deprotonated fraction for titratable residues in CpHMD simulations.

Residues in site A (D47, D51, H155, D159) and site B (D70, H73, H77) from both protomers (A and B) are shown. The instantaneous deprotonated fraction *S* is plotted as a function of simulation time and pH value, sampled across all replicas in the replica exchange simulation. The deprotonated fractions generally converge to a stable value after about 8 ns with similar behavior in protomers A and B, thus indicating sufficient sampling during the CpHMD simulations.

#### Suppl. Fig. 9. Titration curves for titratable residues in CpHMD simulations.

Residues in site A (D47, D51, H155, D159) and site B (D70, H73, H77) are shown from both protomers (A and B). The unprotonated fraction *S* from the end of the CpHMD simulations is shown as a function of pH (black points). The Hill equation (generalized Henderson-Hasselbalch equation) is fitted to the data (black line). The estimated p$K_a$ is indicated as a red line at S=0.5. As described in the text, H73/H77 were considered to be coupled and were analyzed in aggregate; the two p$K_a$s for the combined system (H73AH77A and H73BH77B) are represented by the pair of red lines in the last two panels.

#### Suppl. Fig. 10. Site A Protonation and $Zn^{2+}$-binding states by CpHMD and MST inference.

(A). Populations of protonated states obtained directly from CpHMD simulations for site A, in the absence of $Zn^{2+}$. Data for protomer A and protomer B were combined. Color coded definitions of individual states are shown below with a "1" indicating the protonated state and absence of a number the unprotonated state of each corresponding residue. (B) Populations of protonated states from an inverse *Multibind* model based on inferring microscopic p$K_a$ values directly from the CpHMD populations in (A). (C) Population of protonated states resulting from MST inference, which relies on the MC method to refine microscopic model parameters based on experimental MST data. Populations represent the $Zn^{2+}$-free states although the model incorporates the whole range of $Zn^{2+}$ bound states [see (F)]. Only states S0, S3, S7, S11 have appreciable occupancy (>0.0001 maximum probability) although all states contribute to aggregated state probabilities in D and E. (D) MST inference results represented as populations of aggregated states defined by the total number of protons bound in the absence of $Zn^{2+}$. (E) Deprotonated fraction of each site A residue as a function of pH, averaged over all possible states of the MST inference model in the absence of $Zn^{2+}$. Data points (symbols) were generated from the model; then the Hill-Langmuir equation was fit to the generated data to obtain an effective per-residue p$K_a$ (listed in Table 4). (F) Population of states derived from the MST inference model as a function of $Zn^{2+}$ concentration at various pH values. S0Zn (dotted line) denotes the fully deprotonated, $Zn^{2+}$-bound state, which is the only populated $Zn^{2+}$-bound state predicted by the MST inference.

#### Suppl. Fig. 11. Site B Protonation and $Zn^{2+}$-binding states by CpHMD and MST inference.

(A). Populations of protonated states obtained directly from CpHMD simulations for site B, in the absence of $Zn^{2+}$. Data for protomer A and protomer B were combined. Color coded definitions of individual states are shown below with a "1" indicating the protonated state and absence of a number the unprotonated state of each corresponding residue. (B) Populations of protonated states from an inverse *Multibind* model based on inferring microscopic p$K_a$ values directly from the CpHMD populations in (A). (C) Population of protonated states resulting from MST inference. For this analysis, a symmetry restraint was imposed on H73 and H77 as described in the text. Populations are shown in the absence of $Zn^{2+}$ although the model incorporates the whole range of $Zn^{2+}$ concentrations [see (G)]. (D) MST inference results represented as populations of aggregated states defined by the total number of protons bound in the absence of $Zn^{2+}$. (E) Deprotonated fraction of each site B residue as a function of pH, averaged over all possible states of the MST inference model in the absence of $Zn^{2+}$. Data points were generated from the model and the Hill-Langmuir equation was fit to D70 data (solid black line) to obtain an effective per-residue $pK_a$; data for H73 and H77 are not correctly modelled by a Hill fit (not plotted). (F) H73 and H77 were treated as a coupled system and the total deprotonated fraction as a function of pH was fit with the “coupled titration model” (solid black line), resulting in two effective $pK_a$ values. (G) Population of states from the MST inference mode as a function of $Zn^{2+}$ concentration at various pH values. Unlike site A, multiple $Zn^{2+}$-bound states (S0Zn, S1Zn, S2Zn, S3Zn) are populated at lower pH values.

##### Suppl. Fig. 12. Summary of structures from CDF transporters.

Cartoon representations of most structures so far determined. The structures have been grouped according to conformational state: symmetric IF, symmetric OF and asymmetric states. PDB accession codes are indicated for previously published structures together with concentrations of $Zn^{2+}$ present in the solution and the number of ions observed in the structures. Accession codes for current work are listed in Table 1. Although the sites been named differently for Znt7 and Znt8, the A,B,C nomenclature has been used to simplify the comparison. The helices in the TMD are shown in rainbow colors starting with blue for TM1 and red for TM6. CTD's are represented as a triangle. $Zn^{2+}$ ions as magenta spheres. Hydrophobic residues forming the so-called hydrophobic gate (Leu154 and Leu199 in SoYiiP) are shown as sticks. Disordered in TM2/TM3 loops is indicated by a blurred line.

##### Suppl. Fig. 13. Water accessibility of site A.

Caver Analyst (Jurcik et al., 2018) was used to map cavities starting from Site A and leading towards the cytoplasm. (A) Model of WT YiiP with the cavity depicted a series of spheres, color coded according to the radius. (B) Model of D70A_asym shows an IF conformation on the left and a much constricted cavity in the occluded protomer on the right. (C) Radii are plotted along the contour of the respective cavities shown in panels A and B. This plot shows that the occluded protomer in D70A_asym falls well below the radius of 1.4 Å, typically taken as the limit for water accessibility.

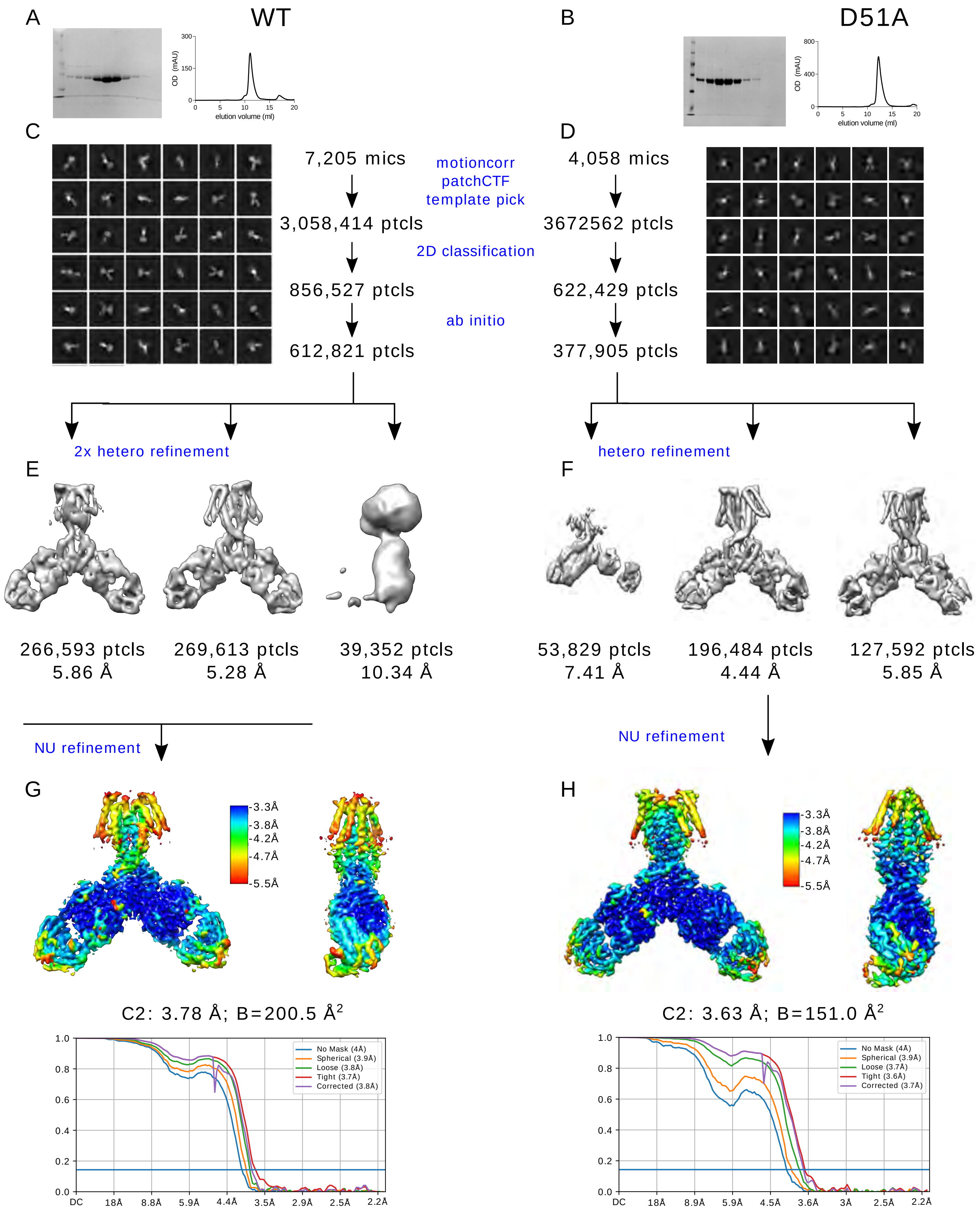

A
WT
B
D51A
OD (mAU)
elution volume (ml)
C
D
7,205 mics
4,058 mics
motioncorr
patchCTF
template pick
3,058,414 ptcls
3672562 ptcls
2D classification
856,527 ptcls
622,429 ptcls
ab initio
612,821 ptcls
377,905 ptcls
2x hetero refinement
hetero refinement
E
F
266,593 ptcls 5.86 Å
269,613 ptcls 5.28 Å
39,352 ptcls 10.34 Å
53,829 ptcls 7.41 Å
196,484 ptcls 4.44 Å
127,592 ptcls 5.85 Å
NU refinement
NU refinement
G
H
-3.3Å
-3.8Å
-4.2Å
-4.7Å
-5.5Å
C2: 3.78 Å; B=200.5 Å2
C2: 3.63 Å; B=151.0 Å2
No Mask (4Å)
Spherical (3.9Å)
Loose (3.8Å)
Tight (3.7Å)
Corrected (3.8Å)
No Mask (4Å)
Spherical (3.9Å)
Loose (3.7Å)
Tight (3.6Å)
Corrected (3.7Å)
DC 18Å 8.8Å 5.9Å 4.4Å 3.5Å 2.9Å 2.5Å 2.2Å
DC 18Å 8.9Å 5.9Å 4.5Å 3.6Å 3Å 2.5Å 2.2Å


Suppl. Fig. 1

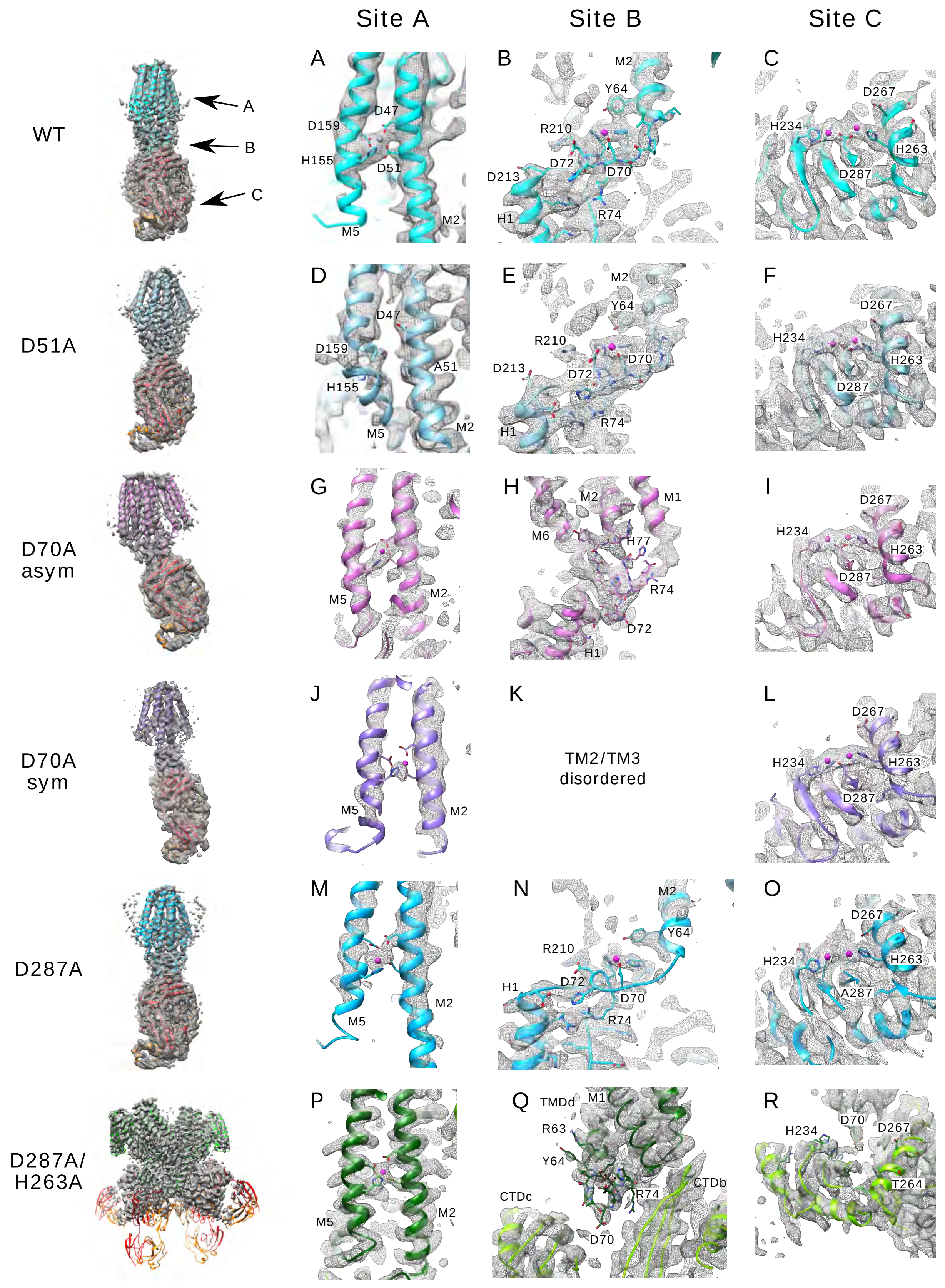
Site A
Site B
Site C
WT
A
B
C
D47
D159
H155
D51
M5
M2
M2
Y64
R210
D72
D70
D213
H1
R74
D267
H234
H263
D287
D51A
D
E
F
D47
D159
A51
H155
M5
M2
M2
Y64
R210
D70
D213
D72
H1
R74
D267
H234
H263
D287
D70A asym
G
H
I
M5
M2
M2
M1
M6
H77
R74
D72
H1
D267
H234
H263
D287
D70A sym
J
K
L
M5
M2
TM2/TM3 disordered
D267
H234
H263
D287
D287A
M
N
O
M5
M2
M2
Y64
R210
H1
D72
D70
R74
D267
H234
H263
A287
D287A/ H263A
P
Q
R
M5
M2
TMDd
M1
R63
Y64
CTDc
R74
CTDb
D70
H234
D70
D267
T264

Suppl. Fig. 2

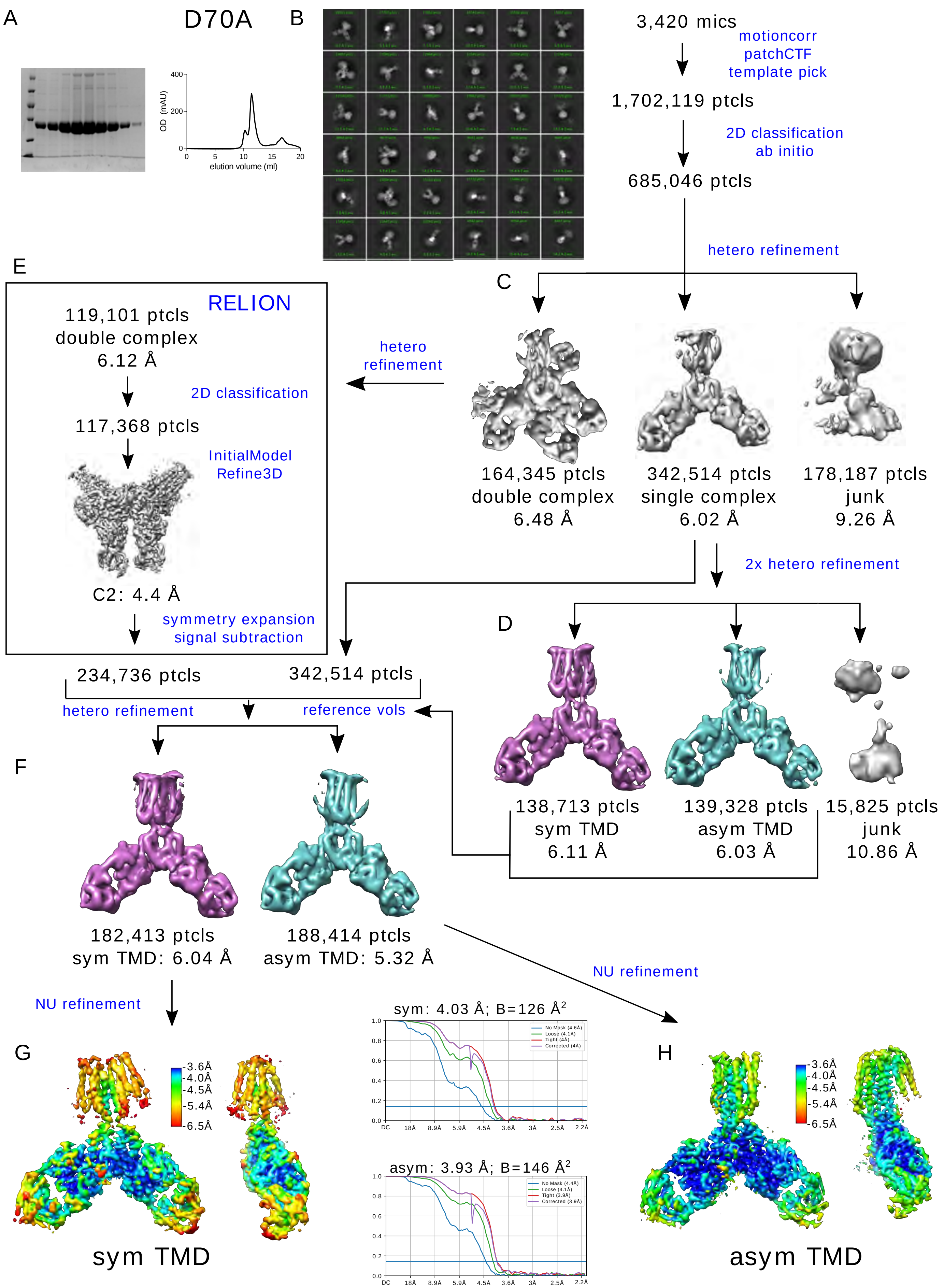
A
D70A
OD (mAU)
400
200
0
0
5
10
15
20
elution volume (ml)
B
3,420 mics
motioncorr
patchCTF
template pick
1,702,119 ptcls
2D classification
ab initio
685,046 ptcls
hetero refinement
C
164,345 ptcls
double complex
6.48 Å
342,514 ptcls
single complex
6.02 Å
178,187 ptcls
junk
9.26 Å
E
RELION
119,101 ptcls
double complex
6.12 Å
2D classification
117,368 ptcls
InitialModel
Refine3D
C2: 4.4 Å
symmetry expansion
signal subtraction
hetero
refinement
2x hetero refinement
D
138,713 ptcls
sym TMD
6.11 Å
139,328 ptcls
asym TMD
6.03 Å
15,825 ptcls
junk
10.86 Å
234,736 ptcls
342,514 ptcls
hetero refinement
reference vols
F
182,413 ptcls
sym TMD: 6.04 Å
188,414 ptcls
asym TMD: 5.32 Å
NU refinement
NU refinement
G
3.6Å
4.0Å
4.5Å
5.4Å
6.5Å
sym TMD
sym: 4.03 Å; B=126 Å$^2$
No Mask (4.6Å)
Loose (4.1Å)
Tight (4Å)
Corrected (4Å)
asym: 3.93 Å; B=146 Å$^2$
No Mask (4.4Å)
Loose (4.1Å)
Tight (3.9Å)
Corrected (3.9Å)
DC
18Å
8.9Å
5.9Å
4.5Å
3.6Å
3Å
2.5Å
2.2Å
H
3.6Å
4.0Å
4.5Å
5.4Å
6.5Å
asym TMD

Suppl. Fig. 3

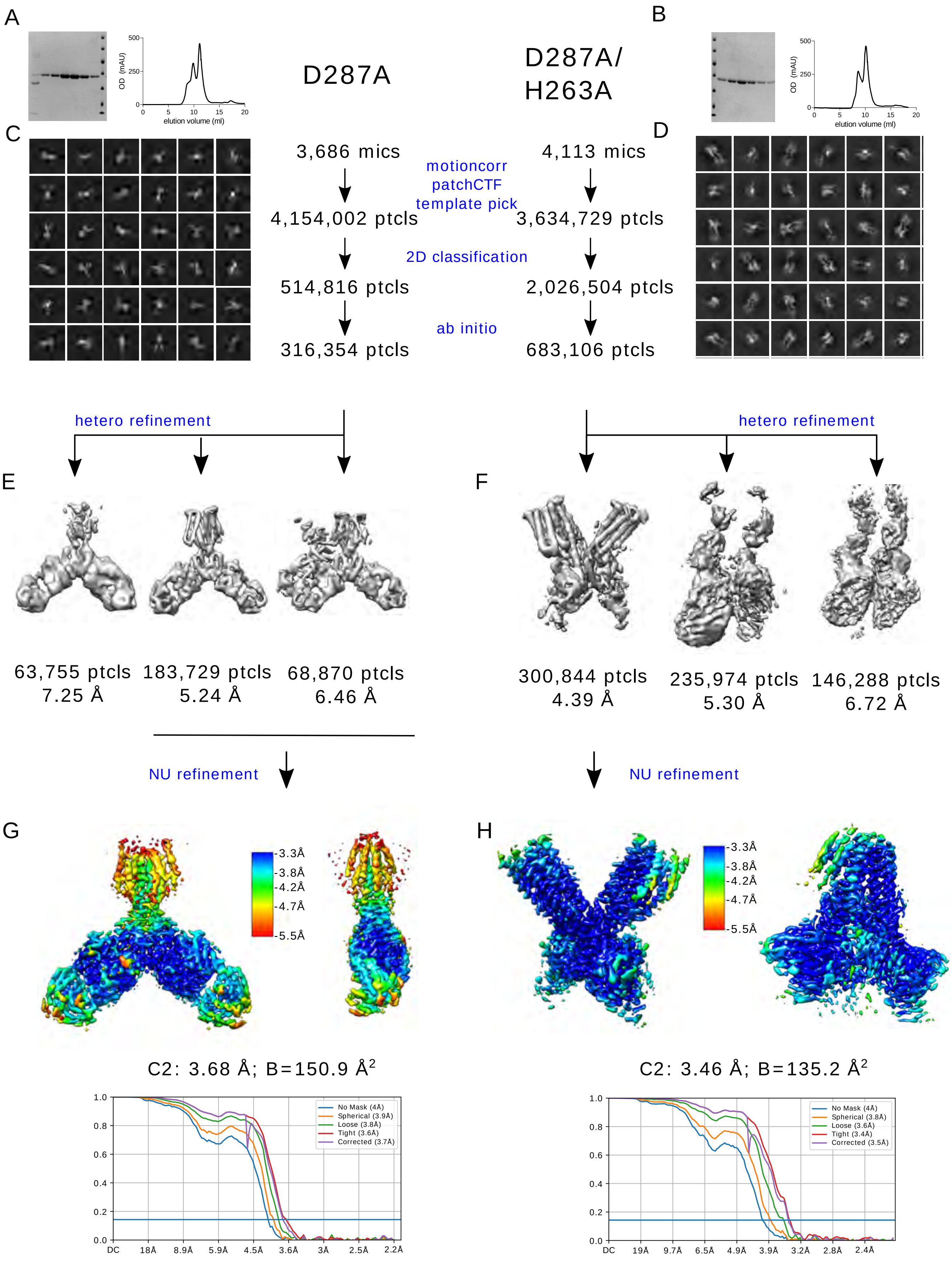
A
B
D287A
D287A/
H263A
OD (mAU)
elution volume (ml)
C
D
3,686 mics
4,113 mics
motioncorr
patchCTF
template pick
4,154,002 ptcls
3,634,729 ptcls
2D classification
514,816 ptcls
2,026,504 ptcls
ab initio
316,354 ptcls
683,106 ptcls
hetero refinement
hetero refinement
E
F
63,755 ptcls
7.25 Å
183,729 ptcls
5.24 Å
68,870 ptcls
6.46 Å
300,844 ptcls
4.39 Å
235,974 ptcls
5.30 Å
146,288 ptcls
6.72 Å
NU refinement
NU refinement
G
H
3.3Å
3.8Å
4.2Å
4.7Å
5.5Å
C2: 3.68 Å; B=150.9 Å2
C2: 3.46 Å; B=135.2 Å2
No Mask (4Å)
Spherical (3.9Å)
Loose (3.8Å)
Tight (3.6Å)
Corrected (3.7Å)
No Mask (4Å)
Spherical (3.8Å)
Loose (3.6Å)
Tight (3.4Å)
Corrected (3.5Å)
DC
18Å
8.9Å
5.9Å
4.5Å
3.6Å
3Å
2.5Å
2.2Å
19Å
9.7Å
6.5Å
4.9Å
3.9Å
3.2Å
2.8Å
2.4Å

Suppl. Fig. 4

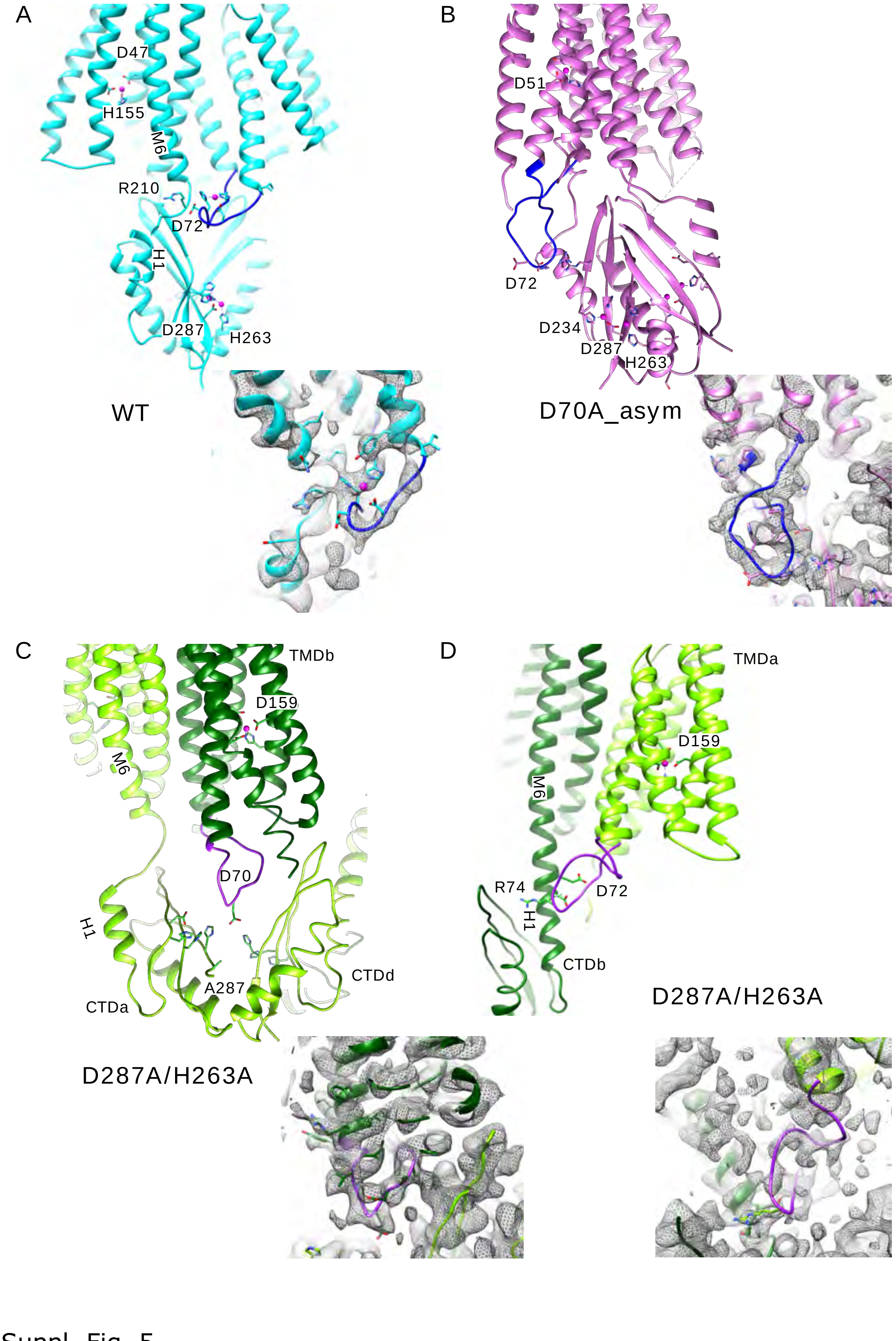
A
D47
H155
M6
R210
D72
H1
D287
H263
WT
B
D51
D72
D234
D287
H263
D70A_asym
C
TMDb
D159
M6
D70
H1
A287
CTDd
CTDa
D287A/H263A
D
TMDa
D159
M6
R74
H1
D72
CTDb
D287A/H263A

Suppl. Fig. 5

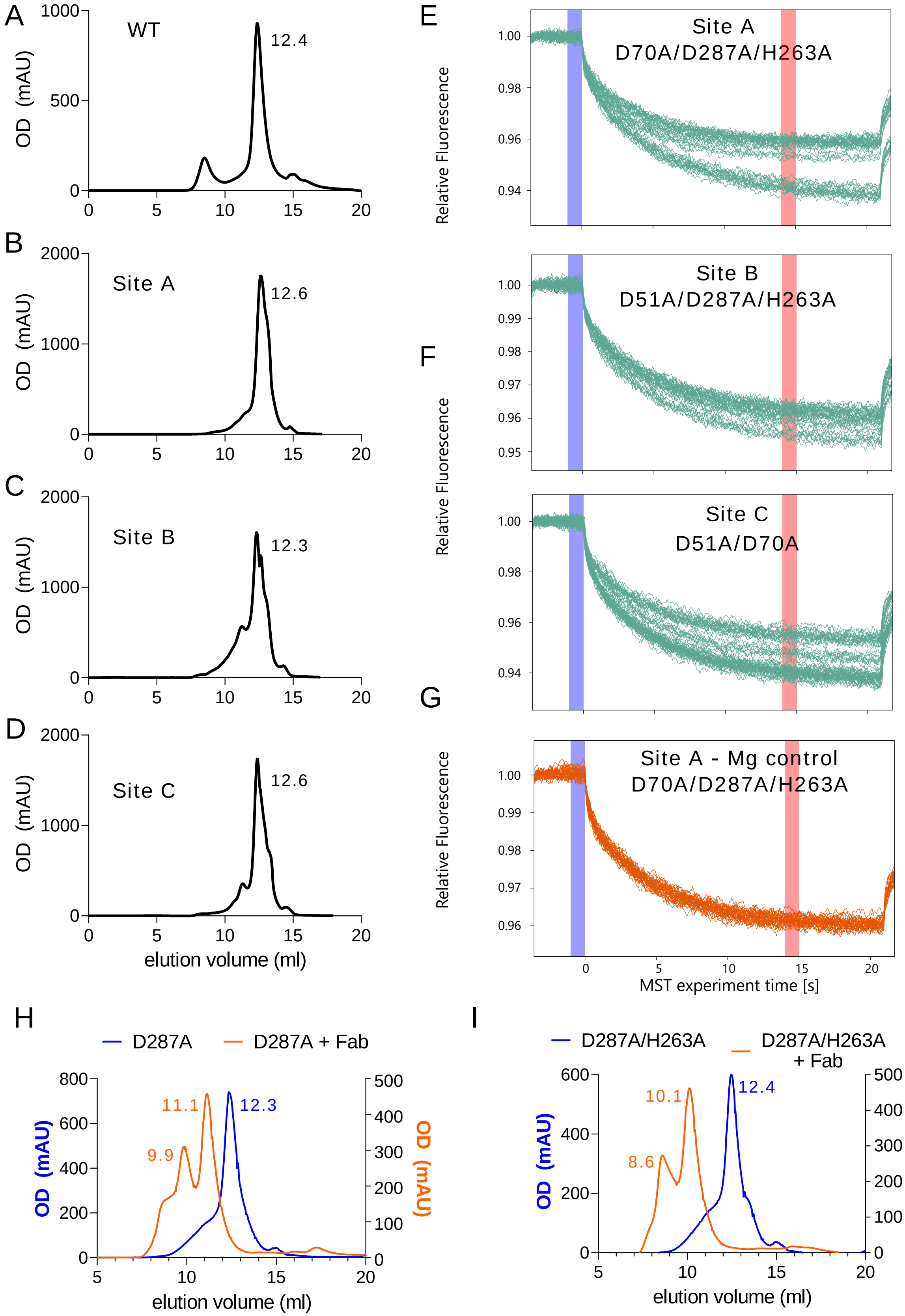
A
WT
12.4
OD (mAU)
B
Site A
12.6
C
Site B
12.3
D
Site C
12.6
elution volume (ml)
E
Site A
D70A/D287A/H263A
Relative Fluorescence
F
Site B
D51A/D287A/H263A
Site C
D51A/D70A
G
Site A - Mg control
D70A/D287A/H263A
MST experiment time [s]
H
D287A
D287A + Fab
11.1
12.3
9.9
I
D287A/H263A
D287A/H263A + Fab
10.1
12.4
8.6

Suppl. Fig. 6

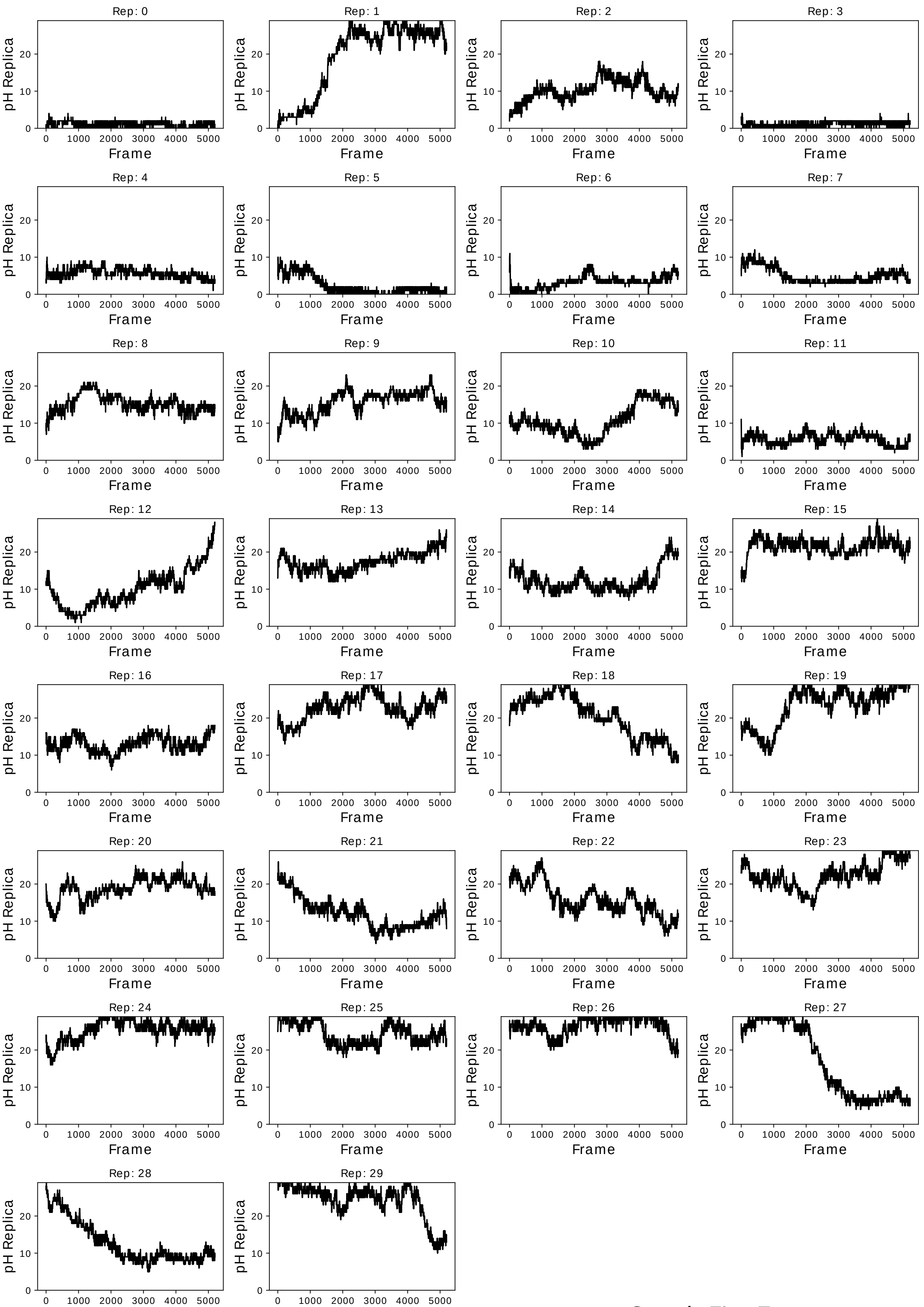
Rep: 0
Rep: 1
Rep: 2
Rep: 3
Rep: 4
Rep: 5
Rep: 6
Rep: 7
Rep: 8
Rep: 9
Rep: 10
Rep: 11
Rep: 12
Rep: 13
Rep: 14
Rep: 15
Rep: 16
Rep: 17
Rep: 18
Rep: 19
Rep: 20
Rep: 21
Rep: 22
Rep: 23
Rep: 24
Rep: 25
Rep: 26
Rep: 27
Rep: 28
Rep: 29
pH Replica
Frame

Suppl. Fig. 7

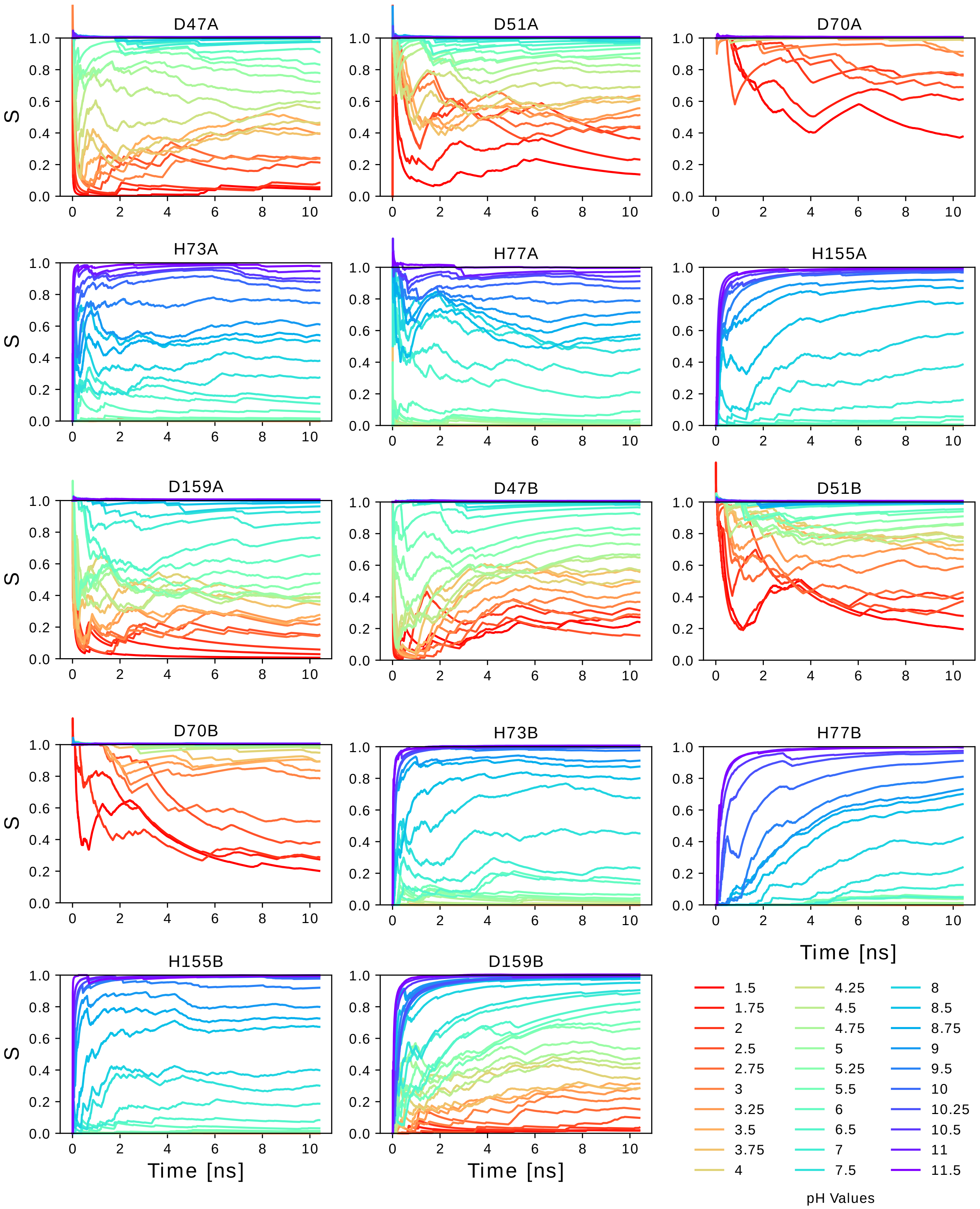
D47A
D51A
D70A
H73A
H77A
H155A
D159A
D47B
D51B
D70B
H73B
H77B
H155B
D159B
S
Time [ns]
pH Values
1.5
1.75
2
2.5
2.75
3
3.25
3.5
3.75
4
4.25
4.5
4.75
5
5.25
5.5
6
6.5
7
7.5
8
8.5
8.75
9
9.5
10
10.25
10.5
11
11.5

Suppl. Fig. 8

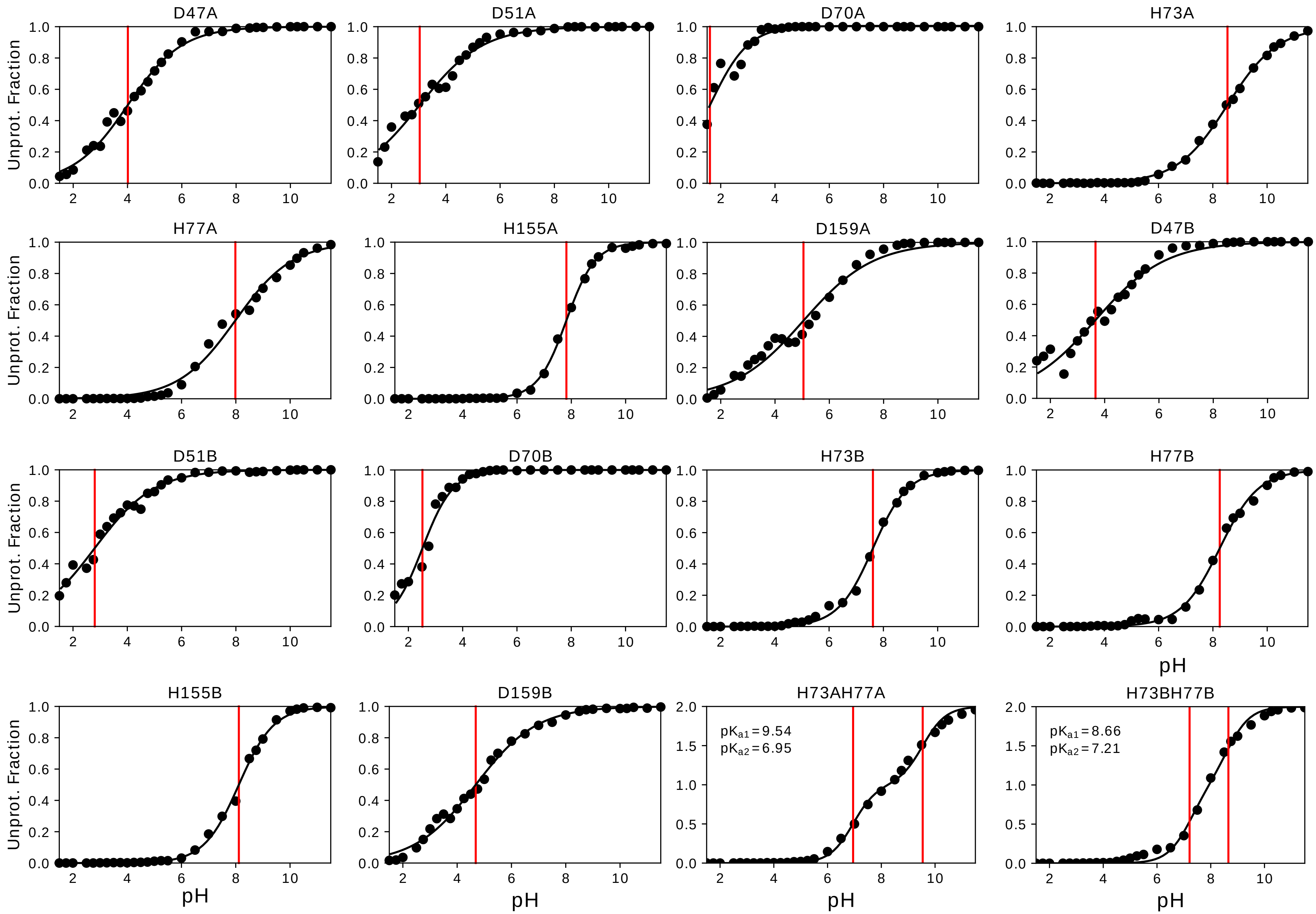
D47A
D51A
D70A
H73A
H77A
H155A
D159A
D47B
D51B
D70B
H73B
H77B
H155B
D159B
H73AH77A
H73BH77B
Unprot. Fraction
pH
$pK_{a1} = 9.54$
$pK_{a2} = 6.95$
$pK_{a1} = 8.66$
$pK_{a2} = 7.21$

Suppl. Fig. 9

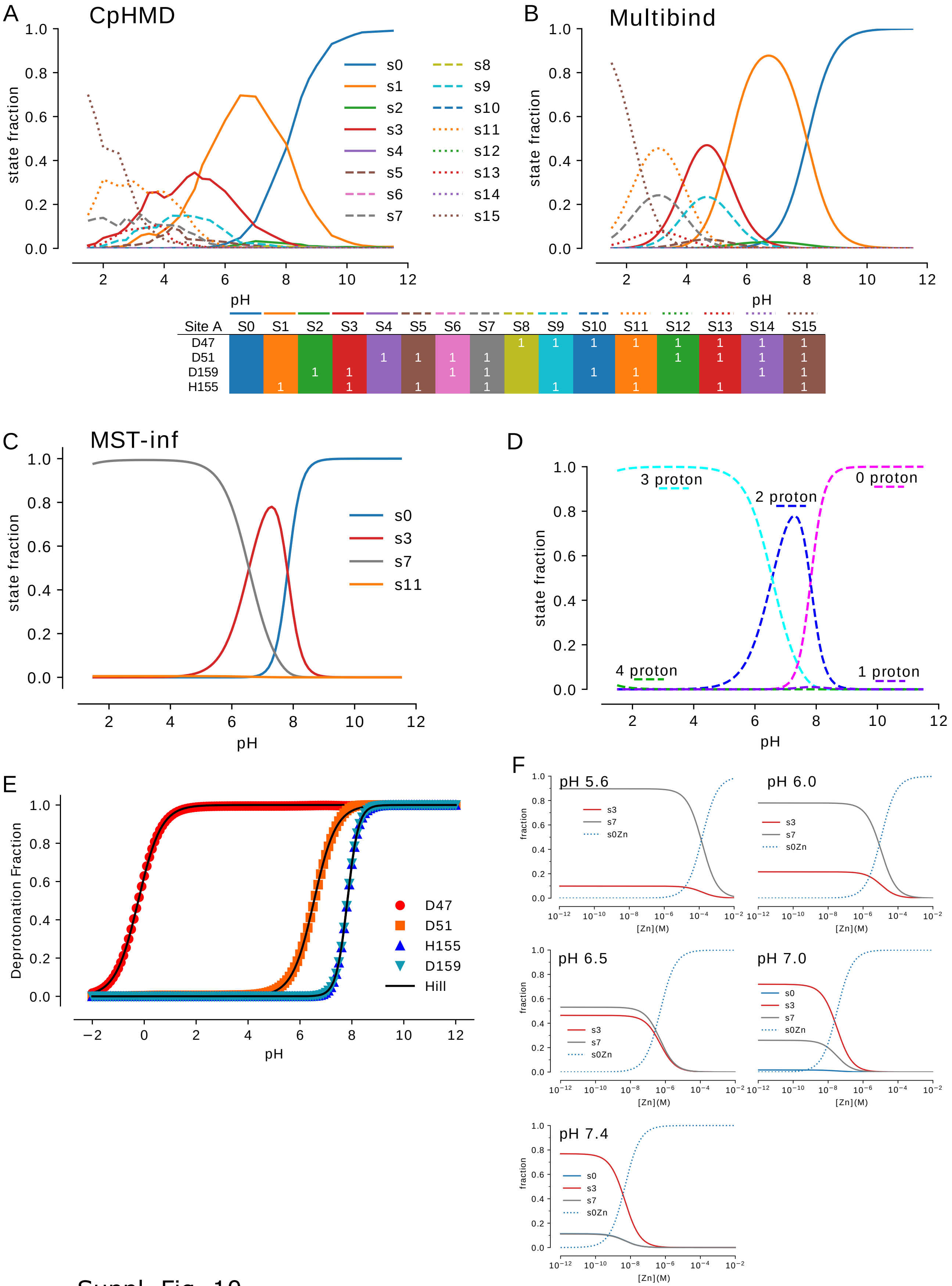


| Site A | S0 | S1 | S2 | S3 | S4 | S5 | S6 | S7 | S8 | S9 | S10 | S11 | S12 | S13 | S14 | S15 |
|---|---|---|---|---|---|---|---|---|---|---|---|---|---|---|---|---|
| D47 | | | | | | | | | 1 | 1 | 1 | 1 | 1 | 1 | 1 | 1 |
| D51 | | | | | 1 | 1 | 1 | 1 | | | | | 1 | 1 | 1 | 1 |
| D159 | | | 1 | 1 | | | 1 | 1 | | | 1 | 1 | | | 1 | 1 |
| H155 | | 1 | | 1 | | 1 | | 1 | | 1 | | 1 | | 1 | | 1 |

Suppl. Fig. 10

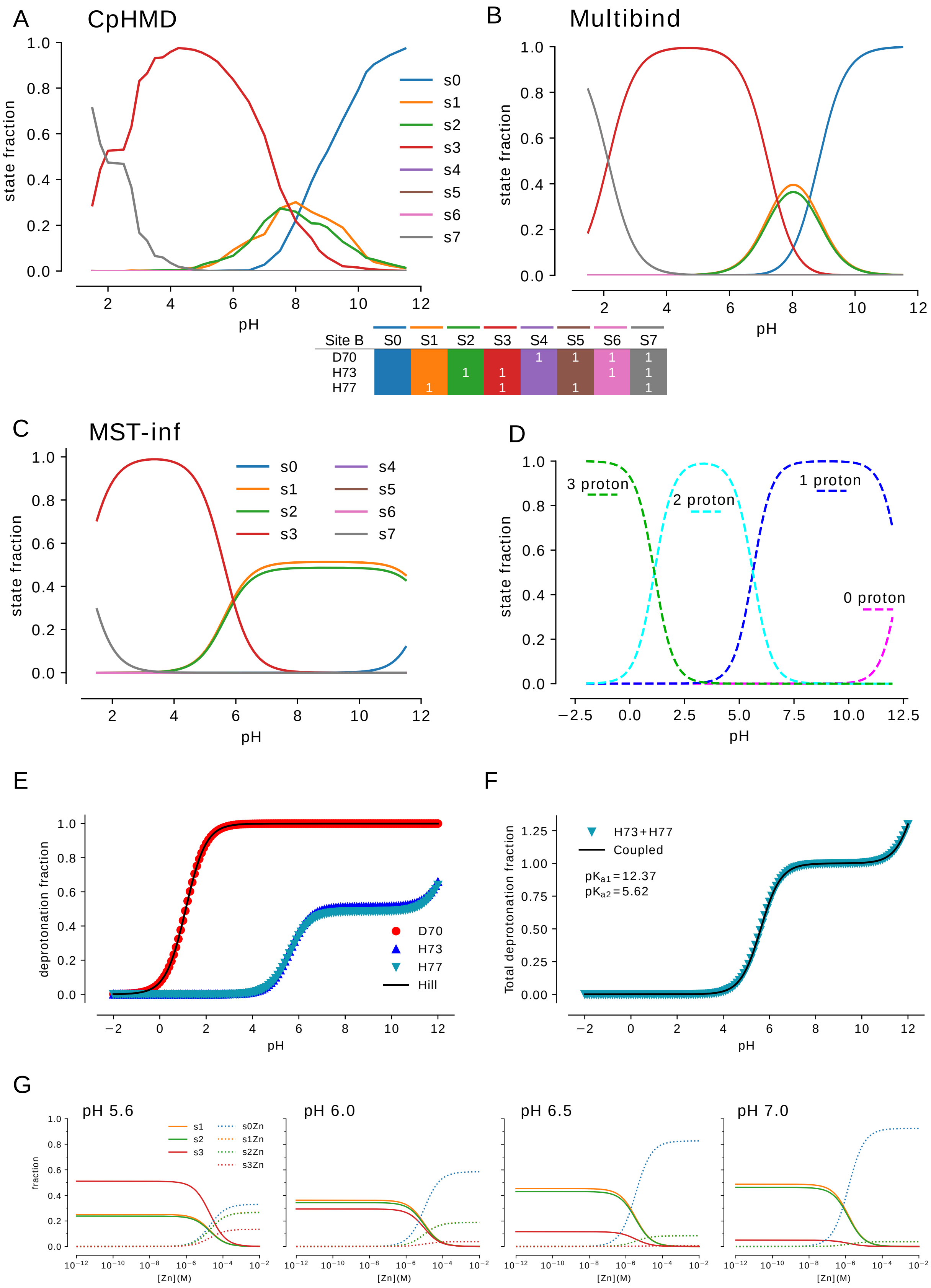

A
CpHMD
B
Multibind
Site B
S0
S1
S2
S3
S4
S5
S6
S7
D70
H73
H77
C
MST-inf
D
3 proton
2 proton
1 proton
0 proton
E
F
H73+H77
Coupled
pKa1=12.37
pKa2=5.62
D70
H73
H77
Hill
G
pH 5.6
pH 6.0
pH 6.5
pH 7.0
state fraction
deprotonation fraction
Total deprotonation fraction
fraction
pH
[Zn](M)


Suppl. Fig. 11

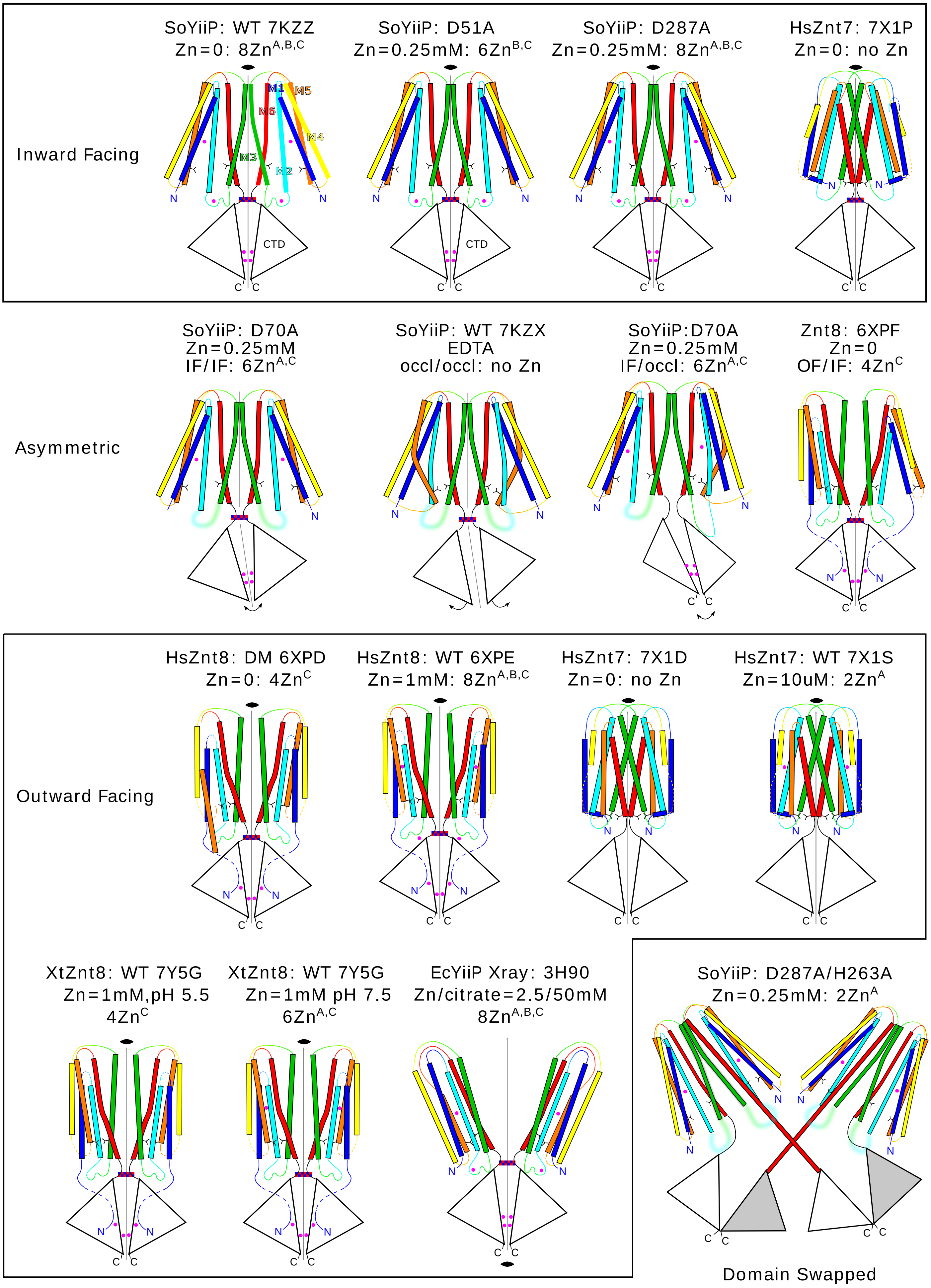

SoYiiP: WT 7KZZ
Zn=0: 8Zn$^{A,B,C}$
SoYiiP: D51A
Zn=0.25mM: 6Zn$^{B,C}$
SoYiiP: D287A
Zn=0.25mM: 8Zn$^{A,B,C}$
HsZnt7: 7X1P
Zn=0: no Zn
Inward Facing
M1
M5
M6
M4
M3
M2
N
CTD
C
SoYiiP: D70A
Zn=0.25mM
IF/IF: 6Zn$^{A,C}$
SoYiiP: WT 7KZX
EDTA
occl/occl: no Zn
SoYiiP:D70A
Zn=0.25mM
IF/occl: 6Zn$^{A,C}$
Znt8: 6XPF
Zn=0
OF/IF: 4Zn$^{C}$
Asymmetric
HsZnt8: DM 6XPD
Zn=0: 4Zn$^{C}$
HsZnt8: WT 6XPE
Zn=1mM: 8Zn$^{A,B,C}$
HsZnt7: 7X1D
Zn=0: no Zn
HsZnt7: WT 7X1S
Zn=10uM: 2Zn$^{A}$
Outward Facing
XtZnt8: WT 7Y5G
Zn=1mM,pH 5.5
4Zn$^{C}$
XtZnt8: WT 7Y5G
Zn=1mM pH 7.5
6Zn$^{A,C}$
EcYiiP Xray: 3H90
Zn/citrate=2.5/50mM
8Zn$^{A,B,C}$
SoYiiP: D287A/H263A
Zn=0.25mM: 2Zn$^{A}$
Domain Swapped


Suppl. Fig. 12

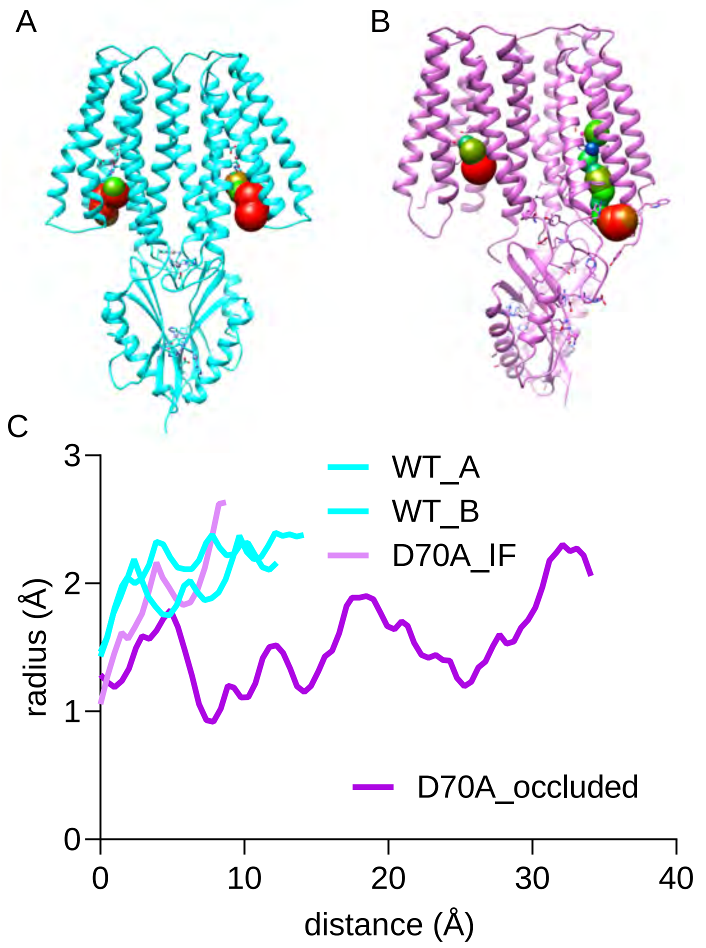
A
B
C
3
2
1
0
radius (Å)
0
10
20
30
40
distance (Å)
WT_A
WT_B
D70A_IF
D70A_occluded

Suppl. Fig. 13
